# Combined Loss of VHL and TSC1 Drives Clear Cell Renal Cell Carcinoma with Metabolic and Redox Reprogramming that Creates an NRF2 Dependency

**DOI:** 10.64898/2026.09.24.754188

**Authors:** Jianing Xu, Xiaoqing Cheng, Omran A. Aboud, Ying-Bei Chen, Ann Bialik, Ed Reznik, Toshinao Oyama, Satish Tickoo, Robert H. Weiss, Emily H. Cheng, James J. Hsieh

**Affiliations:** Human Oncology and Pathogenesis Program, Memorial Sloan Kettering Cancer Center, New York, NY 10065, USA; Molecular Oncology, Department of Medicine, Washington University, St. Louis, MO 63110, USA; Division of Nephrology, University of California Davis, Davis, CA 95616, USA; Department of Pathology and Laboratory Medicine, Memorial Sloan Kettering Cancer Center, New York, NY 10065, USA; Department of Epidemiology and Biostatistics, Memorial Sloan Kettering Cancer Center, New York, NY 10065, USA; Weill Cornell Medicine, New York, NY 10065, USA

**Keywords:** Kidney Cancer, VHL, TSC1, HIF1, mTORC1, NRF2, Cancer Metabolism, Transcriptomics, Metabolomics, Mouse Model

## Abstract

Cancer genomics studies have implicated combined loss of *VHL* and *TSC1* as sufficient to initiate clear cell renal cell carcinoma (ccRCC) in humans. Here, we engineered a mouse model with kidney-specific deletion of *Vhl* and *Tsc1*. *Vhl^F/F^Tsc1^F/F^Ksp-Cre^+^*mice developed multifocal renal cell carcinomas starting at 7 weeks of age with 100% penetrance, and the resulting *Vhl^-/-^Tsc1^-/-^*tumors recapitulated the histopathology of human *VHL^-/-^TSC1^-/-^*ccRCC. Integrated transcriptomic and metabolomic analyses of 4-week-old preneoplastic renal cortices deficient for *Vhl* and *Tsc1* demonstrated heightened HIF and mTORC1 signaling and unexpectedly revealed activation of an antioxidant response regulated by NRF2 (NFE2L2). Biochemical, cell biological, multi-omic, and xenograft studies identified an NRF2 dependency in human *VHL^-/-^TSC1^-/-^*ccRCC cells. Altogether, we report a mouse model that recapitulates human *VHL^-/-^TSC1^-/-^*ccRCC, delineate metabolic and redox reprogramming driven by unrestrained HIF and mTORC1 activation in the renal cortex, and identify NRF2 as a potential therapeutic vulnerability.

---

The number of new kidney cancer cases diagnosed worldwide and in the United States each year is approximately 400,000 and 76,000, respectively^1,2^. Clear cell renal cell carcinoma (ccRCC), the most common and aggressive subtype of kidney cancer (75–80%), features near-universal loss of the *Von Hippel-Lindau* (*VHL*) tumor suppressor gene. *VHL* encodes the substrate-recognition subunit of the VCB (VHL–elongin C–elongin B)–CUL2 E3 ubiquitin ligase complex, which ubiquitinates hypoxia-inducible factor 1α and 2α (HIF1α/HIF2α) for proteasomal degradation^3,4^. Consequently, human ccRCC exhibits conspicuous vascularity due to uncontrolled activation of HIF1α/HIF2α target genes that regulate angiogenesis^5^.

Evidence from patients with germline *VHL* mutations and genetically engineered mouse models with homozygous deletion of *Vhl* demonstrated that loss of VHL alone is insufficient for ccRCC tumorigenesis, indicating the requirement for additional genetic or epigenetic events^6–9^. One candidate pathway is the PI3K/AKT/mTOR pathway, which is altered in approximately 10– 30% of ccRCC^10–12^. Mutations in *TSC1*, *TSC2*, and *MTOR* occur at frequencies of approximately 1.2%, 1%, and 6%, respectively, in TCGA-KIRC and approximately 7%, 3%, and 10%, respectively, in the RECORD-3 and COMPARZ studies (Supplementary Fig. 1a)^13–15^.

Tuberous sclerosis 1 (TSC1; hamartin), together with TSC2 (tuberin) and TBC1D7, forms the TSC complex, which functions as a GTPase-activating protein (GAP) for the small GTPase RHEB and thereby negatively regulates mTOR complex 1 (mTORC1). mTORC1 coordinates cell growth, protein synthesis, and metabolic signaling^16^. One important translational target of mTORC1 is HIF1α, whose protein abundance is further increased upon VHL loss through impaired degradation, thereby enhancing transcription of genes involved in the cellular hypoxic response^17^. Conversely, HIF1α transcriptionally activates REDD1 (DDIT4), which inhibits mTORC1 through the TSC complex, establishing a negative feedback loop that limits HIF1α protein translation^18^.

Elevated levels of reactive oxygen species (ROS) are a common feature of cancer cells^19^. At moderate levels, ROS can function as signaling molecules that promote tumor initiation and progression^20,21^, consistent with the concept of mitohormesis^22^. ROS accumulation in tumors can result from oncogene activation, tumor suppressor loss, altered metabolism, and tumor hypoxia^19,23,24^. However, excessive ROS can cause macromolecular and organelle damage and ultimately lead to cancer cell death^25,26^. Cancer cells therefore activate antioxidant pathways, including the glutathione and thioredoxin systems^27,28^, to maintain redox homeostasis during oncogenic transformation and tumor progression^29^. Notably, our previous metabolomic profiling of 138 paired ccRCC tumors and adjacent normal kidneys demonstrated that tumor aggressiveness is associated with enhanced glutathione metabolism^30^. How redox and glutathione metabolism are regulated in ccRCC remains incompletely understood.

The identification of human *VHL^-/-^TSC1^-/-^*ccRCC suggests that combined loss of *VHL* and *TSC1* may be sufficient to drive ccRCC formation. We therefore generated mice with kidney-specific conditional deletion of *Vhl* and *Tsc1* (hereafter referred to as *Vhl^K-/-^Tsc1^K-/-^*mice). Remarkably, *Vhl^K-/-^Tsc1^K-/-^*mice developed bilateral polycystic kidney disease by 4 weeks of age and multifocal solid renal cell carcinomas within 2 months after birth. Moreover, the histopathology of these mouse tumors recapitulated that of human *VHL^-/-^TSC1^-/-^*ccRCC. To investigate the molecular mechanisms underlying this rapid tumor development, we performed transcriptomic and metabolomic profiling of preneoplastic renal cortices from 4-week-old *Vhl^K-/-^Tsc1^K-/-^*and control mice (*Vhl^K-/-^*, *Tsc1^K-/-^*, and WT), followed by cell-based molecular and biochemical studies and tumor xenograft experiments. Our findings demonstrate that combined loss of VHL and TSC1 drives unrestrained HIF and mTORC1 signaling and extensive metabolic and redox reprogramming, creating an NRF2 dependency that represents a potential therapeutic vulnerability in ccRCC.

## Results

### *Vhl^K-/-^Tsc1^K-/-^*mice display kidney abnormalities and die before 12 weeks of age

To investigate the genetic interaction between *Vhl* and *Tsc1* loss in kidney cancer pathogenesis *in vivo*, we generated mice with kidney-specific conditional deletion of the *Vhl^F/F^* and/or *Tsc1^F/F^*alleles using *Ksp-Cre*, a Cre recombinase driven by the kidney-specific *Cadherin* 16 promoter^31^. The single *Vhl^F/F^Ksp-Cre* and *Tsc1^F/F^Ksp-Cre* conditional knockout mice (hereafter referred to as *Vhl^K-/-^*and *Tsc1^K-/-^*, respectively) have been reported^9,32^. Consistent with prior studies, *Tsc1^K-/-^*mice exhibited early mortality, with none surviving beyond 25 weeks after birth, whereas *Vhl^K-/-^*mice exhibited a slight yet statistically significant decrease in survival (Fig. 1a)^9,32^. Of note, *Vhl^K-/-^Tsc1^K-/-^*mice were born at a frequency of 13.7% (n = 78), lower than the expected Mendelian ratio of 25%. Furthermore, marked postnatal mortality was observed in *Vhl^K-/-^Tsc1^K-/-^*mice, with a maximum lifespan of 12 weeks (Fig. 1a). The incidence of polycystic kidney disease (PKD) was similar between *Tsc1^K-/-^*and *Vhl^K-/-^Tsc1^K-/-^*mice (Fig. 1b), whereas only *Vhl^K-/-^Tsc1^K-/-^*mice displayed hydronephrosis (HN) by 12 weeks of age (Fig. 1c). At 10–11 weeks of age, kidney weights were markedly increased in both *Tsc1^K-/-^*and *Vhl^K-/-^Tsc1^K-/-^* mice compared with WT and *Vhl^K-/-^* mice (Fig. 1d). Serum creatinine levels were elevated in *Tsc1^K-/-^*and *Vhl^K-/-^Tsc1^K-/-^*mice at 4–8 weeks and worsened at 9–12 weeks (Fig. 1e). Representative MRI, gross, histologic, and immunohistochemical examinations of mouse kidneys from all four genotypes are shown in Fig. 1f-i and Supplementary Fig. 1b.

**Figure 1.**
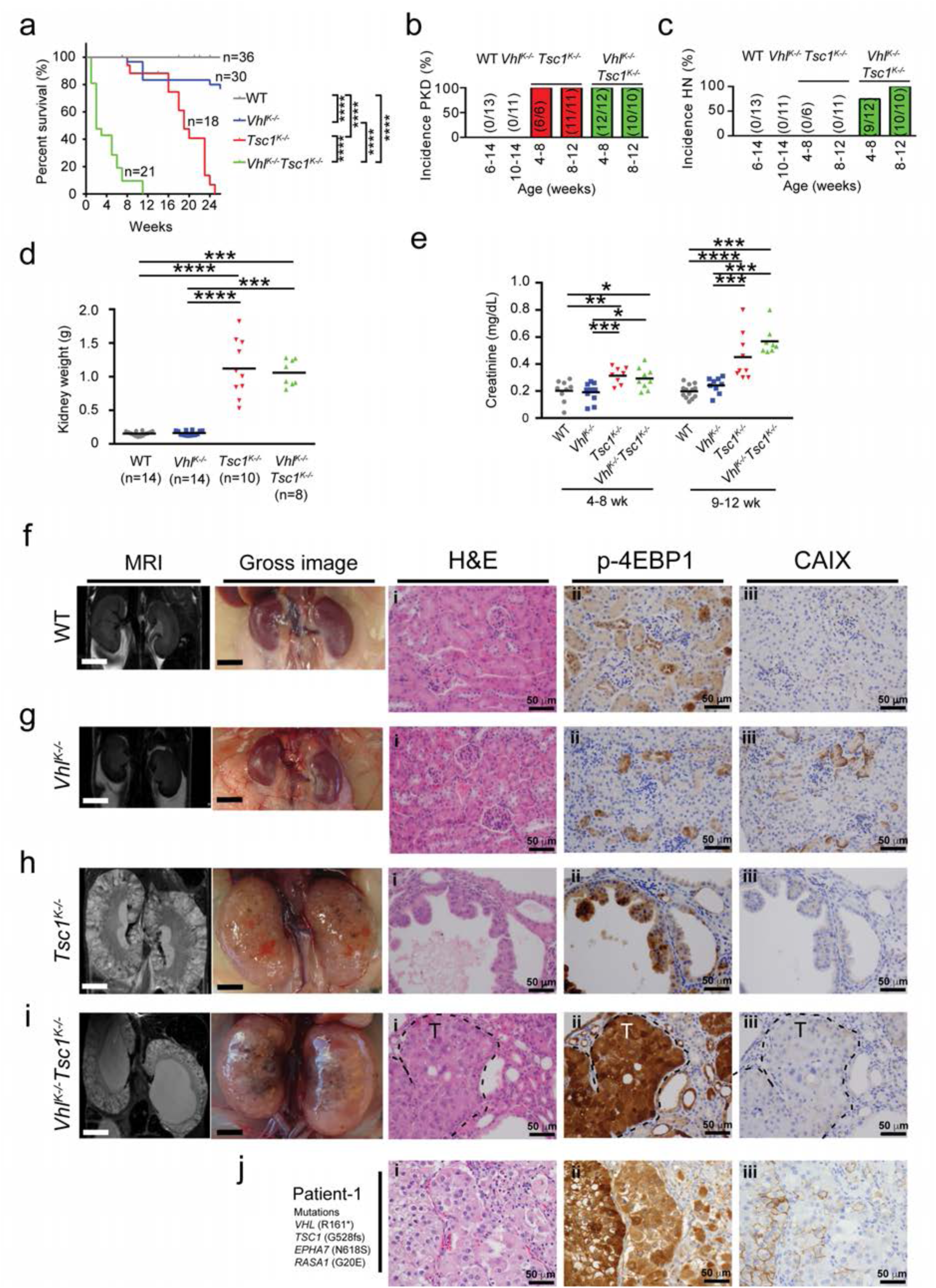
*Vhl^K-/-^Tsc1^K-/-^* mice exhibit increased mortality, kidney abnormalities, and renal cell carcinoma. (a) Kaplan-Meier survival curve of WT, *Vhl^K-/-^, Tsc1^K-/-^,* and *Vhl^K-/-^Tsc1^K-/-^* mice. \*\*\*\**P* < 0.0001 (log-rank (Mantel–Cox) test). (b) Incidence of polycystic kidney disease (PKD) in WT, *Vhl^K-/-^, Tsc1^K-/-^,* and *Vhl^K-/-^Tsc1^K-/-^* mice. The number of animals with PKD relative to the total number of animals in each group is indicated in parentheses. (c) Incidence of hydronephrosis (HN) in WT, *Vhl^K-/-^, Tsc1^K-/-^,* and *Vhl^K-/-^Tsc1^K-/-^* mice. The number of animals with HN relative to the total number of animals in each group is indicated in parentheses. (d) Kidney weights of 10–11-week-old WT, *Vhl^K-/-^, Tsc1^K-/-^,* and *Vhl^K-/-^Tsc1^K-/-^* mice. The number of kidneys analyzed in each group (*n*) is indicated. Data are presented as mean ± SD. \*\*\**P* < 0.001 and \*\*\*\**P* < 0.0001 (Mann–Whitney test). (e) Serum creatinine levels of WT, *Vhl^K-/-^, Tsc1^K-/-^,* and *Vhl^K-/-^Tsc1^K-/-^* mice. Age in weeks is indicated. Data are presented as mean ± SD. \**P* < 0.05, \*\**P* < 0.01, \*\*\**P* < 0.001, and \*\*\*\**P* < 0.0001 (Mann–Whitney test). (f–i) Representative MRI images, gross images, hematoxylin and eosin (H&E)-stained sections (i), and immunohistochemical staining for p-4EBP1 (ii) and CAIX (iii) in kidneys from 10–11-week-old WT (f), *Vhl^K-/-^* (g), *Tsc1^K-/-^* (h), and *Vhl^K-/-^Tsc1^K-/-^*(i) mice. (j) Representative H&E-stained section (i) and immunohistochemical staining for phospho-4EBP1 (ii) and CAIX (iii) of a human renal tumor with *VHL* and *TSC1* loss. The mutation profile determined by MSK-IMPACT analysis is indicated.

### *Vhl^K-/-^Tsc1^K-/-^* mice develop ccRCC that recapitulates the histopathology of human *VHL^-/-^TSC1^-/-^*ccRCC

Upon microscopic examination, *Vhl^K-/-^Tsc1^K-/-^*mice developed multifocal solid renal cell carcinomas starting at ∼7 weeks of age with 100% penetrance by ∼12 weeks (Fig. 1i, Supplementary Fig. 1c, Supplementary Table 1). These *Vhl^K-/-^Tsc1^K-/-^*tumors displayed eosinophilic cytoplasm, nuclear grade 2–3, and increased mitotic activity (Fig. 1i, Supplementary Fig. 1c, Supplementary Table 1). Furthermore, tumor invasion into stromal tissues was observed in large tumors, whereas we did not detect metastasis to the lung, liver, bone, or lymph nodes among 14 tumor-bearing *Vhl^K-/-^Tsc1^K-/-^*mice examined (data not shown). *Vhl^-/-^Tsc1^-/-^* tumors stained positive for lotus tetragonolobus lectin (LTL, a proximal convoluted tubule marker) and negative for Tamm-Horsfall protein (THP, a distal tubule marker) (Supplementary Fig. 1d), consistent with the proximal tubular origin of ccRCC^9,33^. Of note, *Tsc1^K-/-^* mice developed cystadenomas with papillary architecture (Fig. 1h), consistent with a prior report^32^. As expected, both *Vhl^-/-^Tsc1^-/-^*and *Tsc1^-/-^* kidney tumors stained strongly for p-4EBP1 (Fig. 1h,i). *Vhl^-/-^Tsc1^-/-^* tumors were negative for p-AKT and p-ERK but stained strongly for VEGF (Supplementary Fig. 1e). Positive staining for the endothelial cell marker CD31 was detected at the tumor periphery (Supplementary Fig. 1e). Compared with adjacent non-tumor kidney epithelial cells, *Vhl^-/-^Tsc1^-/-^* tumor cells displayed increased proliferation, as determined by Ki-67 staining, but no increase in apoptotic cell death, as assessed by cleaved caspase-3 and TUNEL staining (Supplementary Fig. 1f). Notably, similar to human *VHL^-/-^TSC1^-/-^*ccRCC, mouse *Vhl^-/-^Tsc1^-/-^* kidney tumors were mostly negative for CAIX (Fig. 1i,j, Supplementary Fig. 1g, Supplementary Table 2), in stark contrast to the strong CAIX staining observed in common human ccRCC subtypes such as *VHL^-/-^ PBRM1^-/-^* and in mouse *Vhl^-/-^Pbrm1^-/-^* ccRCC^9^.

### Combined loss of VHL and TSC1 enhances HIF and mTORC1 transcriptional outputs in 4-week-old preneoplastic renal cortices

To gain insight into the transcriptional mechanisms underlying the rapid development of ccRCC following combined deletion of *Vhl* and *Tsc1*, we performed RNA-seq on renal cortices from 4-week-old mice, a stage at which no microscopic tumors are detectable, thereby minimizing potential confounding by tumor mass (*n* = 4 per genotype) (Supplementary Table 5). Principal component analysis (PCA) showed that samples of the same genotype clustered together (Supplementary Fig. 2a). WT and *Vhl^K-/-^* cortices clustered in close proximity, whereas *Tsc1^K-/-^* and *Vhl^K-/-^Tsc1^K-/-^*cortices formed distinct and well-separated clusters. Unsupervised hierarchical clustering of genes differentially expressed in at least one genotype identified three major clusters (Fig. 2a). Clusters I and II comprised genes uniquely downregulated and upregulated, respectively, in *Vhl^K-/-^Tsc1^K-/-^*cortices, whereas Cluster III contained genes upregulated in both *Vhl^K-/-^Tsc1^K-/-^*and *Tsc1^K-/-^* cortices (Fig. 2a). Mechanistically, Cluster I was enriched for mitochondrial and oxidative phosphorylation (OXPHOS) genes, Cluster II for angiogenesis and HIF target genes, and Cluster III for ribosome, lysosome, and mTORC1 pathway genes (Fig. 2b and Supplementary Table 3). Compared with either *Vhl^K-/-^* or *Tsc1^K-/-^*cortices, *Vhl^K-/-^Tsc1^K-/-^* cortices exhibited more profound repression of OXPHOS genes and stronger activation of hypoxia and mTORC1 signature genes, suggesting that concurrent activation of HIF and mTORC1 establishes a positive feedforward circuit that amplifies the transcriptional outputs of both pathways (Fig. 2c–e and Supplementary Table 4). Further supporting enhanced HIF transcriptional activity, qRT-PCR analysis confirmed the expected regulation of key HIF-responsive genes, including upregulation of *Pdk1* (pyruvate dehydrogenase kinase 1) and *Ddit4* (DNA damage-inducible transcript 4) and downregulation of *Ndufa2* (NADH:ubiquinone oxidoreductase subunit A2) and *Sdhd* (succinate dehydrogenase subunit D) in *Vhl^K-/-^Tsc1^K-/-^* cortices (Fig. 2f). Of note, previous studies have shown that HIF indirectly suppresses PGC1α and PGC1β through activation of DEC1 and inhibition of MYC, respectively^34,35^. Furthermore, immunohistochemical analysis of kidneys from 11-week-old WT, *Vhl^K-/-^*, *Tsc1^K-/-^*, and *Vhl^K-/-^Tsc1^K-/-^*mice revealed robust accumulation of HIF1α protein in *Vhl^K-/-^Tsc1^K-/-^* tumors (Supplementary Fig. 2b). These findings indicate that combined loss of VHL and TSC1 is associated with increased HIF1α protein abundance and enhanced HIF transcriptional activity (Fig. 2d).

**Figure 2.**
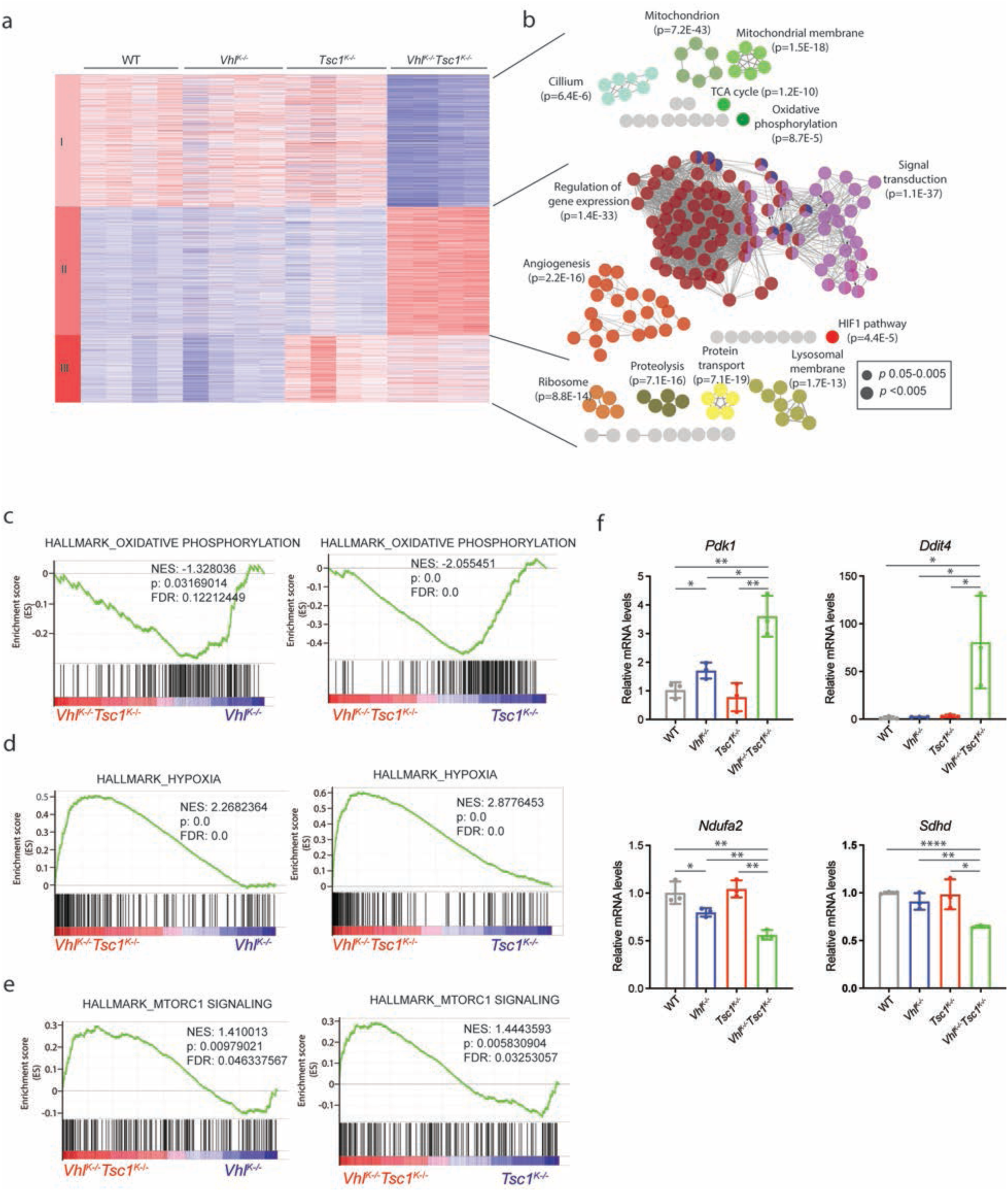
Combined loss of VHL and TSC1 enhances HIF and mTORC1 transcriptional outputs in preneoplastic renal cortices. (a) Unsupervised hierarchical clustering of differentially expressed genes in renal cortices from 4-week-old WT, *Vhl^K-/-^, Tsc1^K-/-^,* and *Vhl^K-/-^Tsc1^K-/-^* mice (*n* = 4 per genotype; Benjamini–Hochberg-adjusted *P* value ≤ 0.05). Clusters I–III are indicated. (b) ClueGO enrichment analysis of genes in Clusters I–III identified in (a). Functionally related enriched pathways and biological processes are shown as networks. (c–e) Gene set enrichment analysis (GSEA) comparing *Vhl^K-/-^Tsc1^K-/-^* renal cortices with *Vhl^K-/-^* or *Tsc1^K-/-^* renal cortices using the MSigDB HALLMARK_OXIDATIVE_PHOSPHORYLATION (c), HALLMARK_HYPOXIA (d), and HALLMARK_MTORC1_SIGNALING (e) gene sets. (f) Relative mRNA levels of the HIF-responsive genes *Pdk1*, *Ddit4*, *Ndufa2*, and *Sdhd* in renal cortices from 4-week-old WT, *Vhl^K-/-^, Tsc1^K-/-^,* and *Vhl^K-/-^Tsc1^K-/-^* mice, measured by qRT-PCR. Data are presented as mean ± SD. \**P* < 0.05, \*\**P* < 0.01, \*\*\**P* < 0.001, and \*\*\*\**P* < 0.0001 (Student’s *t*-test).

### Metabolomic analysis reveals extensive metabolic rewiring in *Vhl^K-/-^Tsc1^K-/-^* renal cortices

As HIF and mTORC1 are major regulators of cellular metabolism, we directly assessed global metabolic alterations in *Vhl^K-/-^Tsc1^K-/-^*renal cortices using gas chromatography time-of-flight mass spectrometry (GC-TOF/MS)^36^. Freshly dissected renal cortices from 4-week-old mice were snap-frozen for metabolomic analysis (n = 6 per genotype; 24 samples in total). Principal component analysis (PCA) showed that all *Vhl^K-/-^Tsc1^K-/-^*samples clustered together and were clearly separated from the other genotypes (Fig. 3a). One-way ANOVA identified 113 metabolites that differed significantly in abundance among the four genotypes (WT*, Vhl^K-/-^, Tsc1^K-/-^, and Vhl^K-/-^Tsc1^K-/-^*), with an FDR-adjusted *P* value ≤ 0.05 (Supplementary Table 6). *Vhl^K-/-^Tsc1^K-/-^*cortices exhibited the greatest number of uniquely altered metabolites (Fig. 3b). To further characterize these metabolic alterations, we performed pathway analyses comparing *Vhl^K-/-^Tsc1^K-/-^* cortices with WT, *Vhl^K-/-^*, and *Tsc1^K-/-^* cortices (Fig. 3c). These analyses revealed extensive metabolic rewiring, with alterations in aspartate metabolism, ammonia recycling, the urea cycle, and β-alanine, tryptophan, tyrosine, purine, glutamate, nicotinate and nicotinamide, glutathione, arginine, and proline metabolism shared across at least two comparisons.

**Figure 3.**
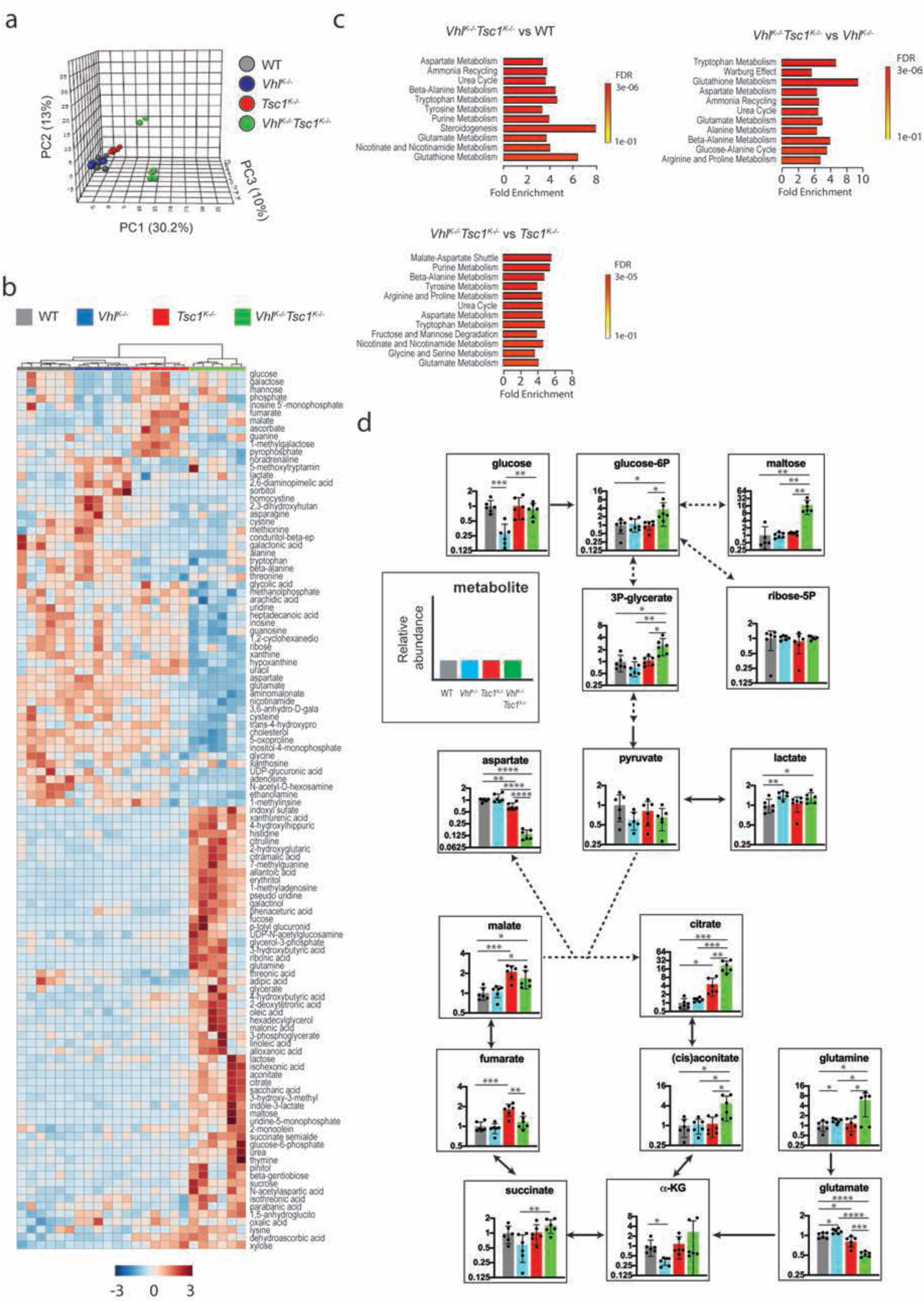
Metabolomic profiling reveals extensive metabolic reprogramming in *Vhl^K-/-^Tsc1^K-/-^* renal cortices. (a) Principal component analysis (PCA) of metabolomic profiles of renal cortices from 4-week-old WT, *Vhl^K-/-^, Tsc1^K-/-^,* and *Vhl^K-/-^Tsc1^K-/-^* mice. (b) Heat map of normalized abundances of metabolites that differed significantly among renal cortices from 4-week-old WT, *Vhl^K-/-^, Tsc1^K-/-^,* and *Vhl^K-/-^Tsc1^K-/-^* mice (FDR-adjusted *P* value ≤ 0.05). (c) Quantitative metabolic pathway enrichment analysis using MetaboAnalyst comparing *Vhl^K-/-^ Tsc1^K-/-^* with WT, *Vhl^K-/-^*, or *Tsc1^K-/-^* renal cortices. Pathways with FDR < 0.1 are shown. (d) Relative abundances of measured metabolites involved in central carbon metabolism in renal cortices from 4-week-old mice of the indicated genotypes. Data are presented as mean ± SD. \**P* < 0.05, \*\**P* < 0.01, \*\*\**P* < 0.001, and \*\*\*\**P* < 0.0001 (Student’s *t*-test).

A key aspect of metabolic reprogramming during tumorigenesis involves the utilization of carbon for biosynthesis and energy production (Supplementary Fig. 3a)^37^. Remarkably, when metabolite levels across the four genotypes were mapped onto major cellular carbon metabolic pathways, *Vhl^K-/-^Tsc1^K-/-^* renal cortices exhibited increased glycogen synthesis and glycolysis (Fig. 3d), consistent with the transcriptomic data (Supplementary Fig. 3b,c). Notably, despite marked downregulation of genes involved in the TCA cycle and mitochondrial biogenesis (Supplementary Fig. 3d,e), several TCA cycle metabolites, including citrate, cis-aconitate, and malate, were elevated in *Vhl^K-/-^Tsc1^K-/-^*cortices (Fig. 3d). A similar discordance between transcriptomic and metabolomic alterations was observed in our previous global metabolomic analysis of human ccRCC^30^, highlighting the value of an integrated multi-omics approach to studying kidney cancer.

Of note, a recent study demonstrated that fumarate was the most highly elevated metabolite in *Tsc1^K-/-^* kidneys and that the resulting protein succination contributed to oncogenesis^32^. Consistent with this report, we observed increased fumarate in *Tsc1^K-/-^* renal cortices; however, no increase in fumarate was detected in *Vhl^K-/-^Tsc1^K-/-^* cortices (Supplementary Fig. 3f). Furthermore, human *VHL^-/-^TSC1^-/-^* ccRCC showed negative staining for S-(2-succinyl)cysteine (2SC), a surrogate marker of fumarate accumulation (Supplementary Fig. 3g). Together, these findings suggest that fumarate is unlikely to be a major oncometabolite contributing to ccRCC tumorigenesis in either mouse *Vhl^-/-^Tsc1^-/-^* or human *VHL^-/-^TSC1^-/-^*ccRCC.

### Mouse *Vhl^-/-^Tsc1^-/-^* kidney tumors exhibit heightened redox and NRF2 signaling

Notably, our previous global metabolomic study of human kidney cancer, which analyzed 877 metabolites in 138 paired ccRCC tumors and adjacent normal kidneys, identified four metabolic clusters, among which the cluster associated with the poorest clinical outcome exhibited enhanced glutathione and cysteine metabolism^30^. Therefore, among the pathway alterations identified by our integrated transcriptomic and metabolomic analyses of *Vhl^K-/-^Tsc1^K-/-^* mouse renal cortices, we focused on the prominent redox reprogramming characterized by enhanced glutathione metabolism (Fig. 3c).

As a major cellular antioxidant, glutathione rapidly interconverts between its reduced (GSH) and oxidized (GSSG) forms to buffer oxidative stress, and the GSH/GSSG ratio reflects cellular reducing capacity (Fig. 4a)^27^. We first measured the GSH/GSSG ratio in renal cortices. The ratio was highest in *Vhl^K-/-^Tsc1^K-/-^* cortices among the four genotypes, reaching approximately twofold the level observed in WT or *Vhl^K-/-^* cortices (Fig. 4b). Notably, GSH levels did not differ significantly among the genotypes, whereas GSSG levels were significantly reduced in *Vhl^K-/-^ Tsc1^K-/-^* cortices (Fig. 4b). Glutathione is synthesized through two sequential reactions catalyzed by glutamate-cysteine ligase (GCL) and glutathione synthetase (GSS), both of which are regulated downstream of NRF2^38,39^. Remarkably, GCLC, the catalytic subunit of GCL, was increased approximately fourfold in *Vhl^K-/-^Tsc1^K-/-^*renal cortices (Fig. 4c), suggesting activation of NRF2 signaling. Consistent with this, *Nfe2l2* (NRF2) mRNA levels were approximately twofold higher in *Vhl^K-/-^Tsc1^K-/-^*cortices than in the other genotypes (Fig. 4c).

**Figure 4.**
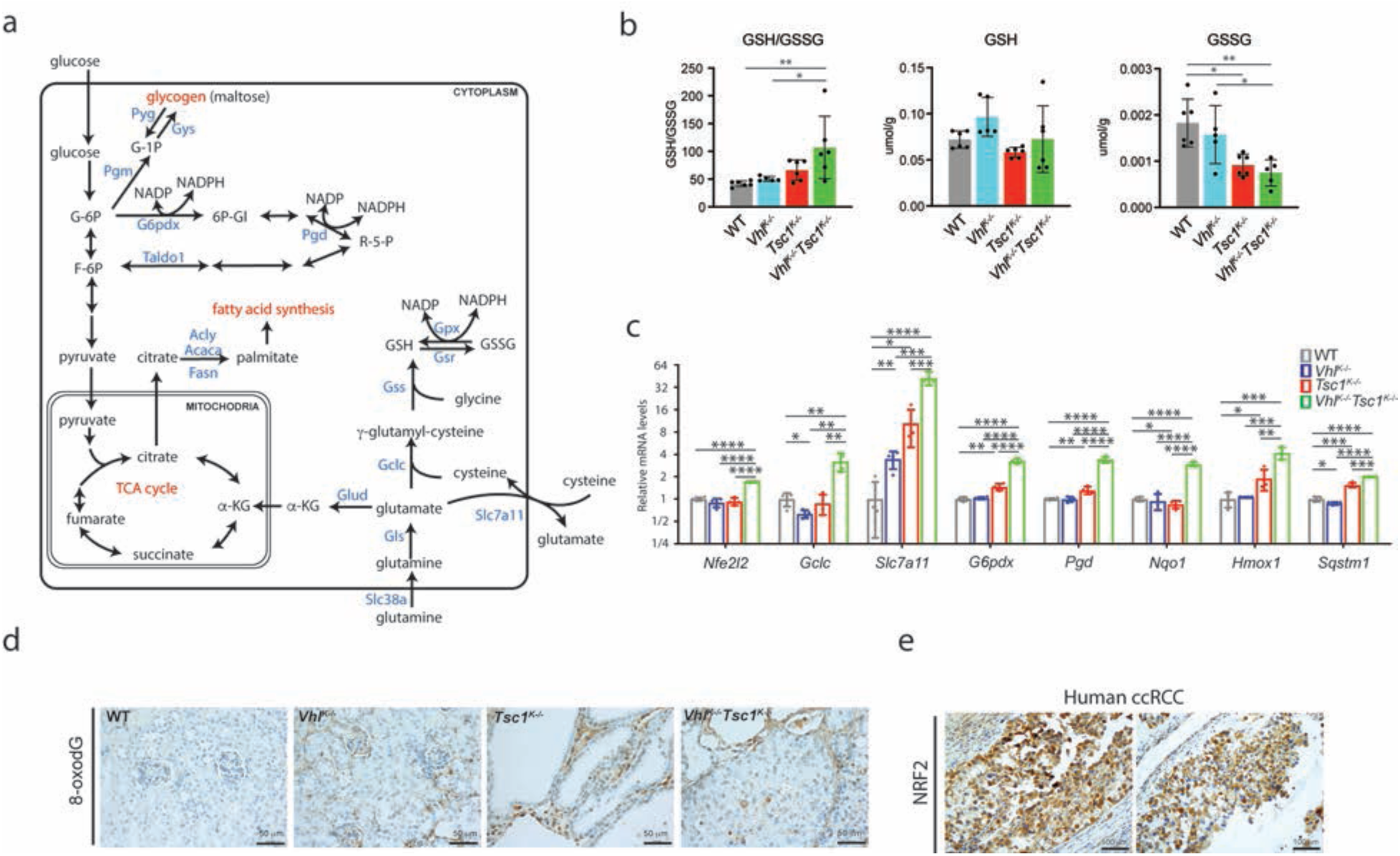
Combined loss of VHL and TSC1 enhances NRF2-dependent antioxidant signaling. (a) Schematic of central carbon and glutathione metabolism. (b) GSH/GSSG ratios and abundances of GSH and GSSG in renal cortices from 4-week-old mice of the indicated genotypes. (c) Relative mRNA levels of *Nfe2l2* (NRF2) and NRF2 target genes in renal cortices from 4-week-old mice of the indicated genotypes. (d) Representative immunohistochemical staining for 8-oxo-dG in kidneys from 11-week-old WT, *Vhl^K-/-^, Tsc1^K-/-^,* and *Vhl^K-/-^Tsc1^K-/-^* mice. (e) Representative immunohistochemical staining for NRF2 in human *VHL^-/-^TSC1^-/-^* ccRCC tumors from patients 4 and 5 in Supplementary Fig. 1g. Data in (b) and (c) are presented as mean ± SD. \**P* < 0.05, \*\**P* < 0.01, \*\*\**P* < 0.001, and \*\*\*\**P* < 0.0001 (Student’s *t*-test).

NRF2, a master regulator of cellular redox homeostasis^40,41^, also controls the expression of multiple antioxidant and redox-related genes, including *Slc7a11*, *G6pdx*, *Pgd*, *Nqo1*, *Hmox1*, and *Sqstm1*^38^. We therefore examined these NRF2 target genes in mouse renal cortices and found that all were significantly upregulated in *Vhl^K-/-^Tsc1^K-/-^*cortices compared with the other genotypes (Fig. 4c). We reasoned that this enhanced antioxidant program might mitigate oxidative stress in *Vhl^-/-^Tsc1^-/-^* kidney tumors. Consistent with this possibility, immunohistochemical staining for 8-oxo-dG, a marker of oxidative DNA damage^42^, did not reveal increased DNA oxidation in *Vhl^-/-^ Tsc1^-/-^* tumors (Fig. 4d). Moreover, consistent with our findings in mouse *Vhl^-/-^Tsc1^-/-^* tumors, NRF2 expression was also elevated in human *VHL^-/-^TSC1^-/-^* ccRCC (Fig. 4e).

### Hyperactive mTORC1 signaling promotes NRF2 protein expression and activity in human ccRCC

The increased NRF2 signaling observed in *Vhl^K-/-^Tsc1^K-/-^*mouse renal cortices suggests that, in the context of *Vhl* deficiency, NRF2 may be upregulated in part through unrestrained mTORC1 signaling. Indeed, NRF2 and mTORC1 signaling showed a strong positive correlation in both the TCGA ccRCC (KIRC) dataset and the CCLE kidney cancer cell-line dataset (Fig. 5a). Consistent with the established relationship between oxidative stress and NRF2 activation, an even stronger positive correlation between NRF2 and reactive oxygen species (ROS) signaling was observed in both datasets (Fig. 5b).

**Figure 5.**
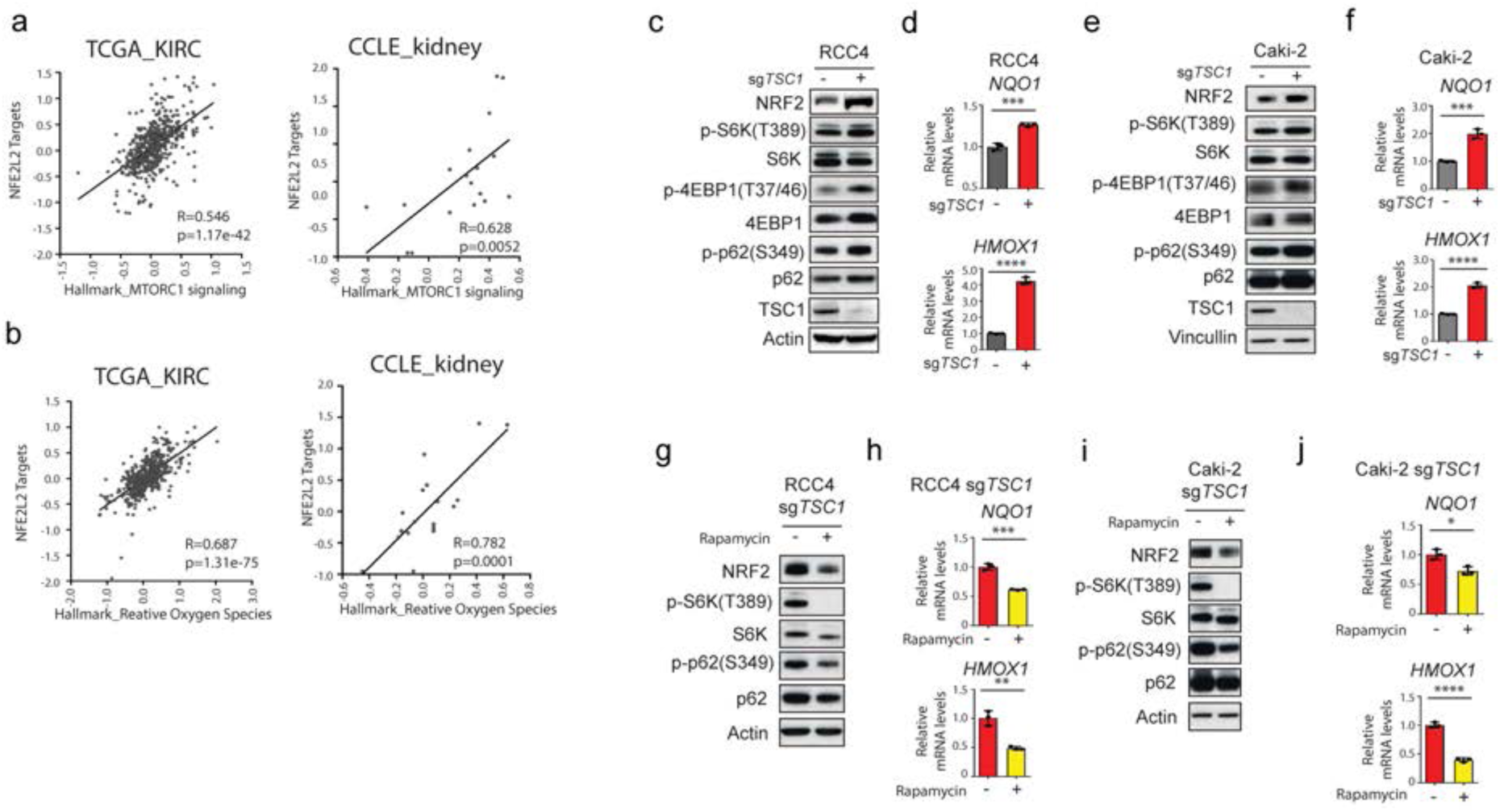
Hyperactive mTORC1 signaling promotes NRF2 expression and activity in human ccRCC. (a) Scatter plots showing the correlation between the SINGH_NFE2L2_TARGETS and HALLMARK_MTORC1_SIGNALING gene signatures in the TCGA-KIRC dataset and the kidney cancer subset of the CCLE dataset. (b) Scatter plots showing the correlation between the SINGH_NFE2L2_TARGETS and HALLMARK_REACTIVE_OXYGEN_SPECIES gene signatures in the TCGA-KIRC dataset and the kidney cancer subset of the CCLE dataset. (c) RCC4 cells expressing sg*LacZ* (−) or sg*TSC1* (+) were analyzed by immunoblotting with the indicated antibodies. (d) Relative mRNA levels of *NQO1* and *HMOX1* in RCC4 cells expressing sg*LacZ* (−) or sg*TSC1* (+), measured by qRT-PCR. (e) Caki-2 cells expressing sg*LacZ* (−) or sg*TSC1* (+) were analyzed by immunoblotting with the indicated antibodies. (f) Relative mRNA levels of *NQO1* and *HMOX1* in Caki-2 cells expressing sg*LacZ* (−) or sg*TSC1* (+), measured by qRT-PCR. (g) RCC4 sg*TSC1* cells were treated with DMSO (−) or 25 nM rapamycin (+) overnight and analyzed by immunoblotting with the indicated antibodies. (h) Relative mRNA levels of *NQO1* and *HMOX1* in RCC4 sg*TSC1* cells treated with DMSO (−) or 25 nM rapamycin (+) overnight, measured by qRT-PCR. (i) Caki-2 sg*TSC1* cells were treated with DMSO (−) or 25 nM rapamycin (+) overnight and analyzed by immunoblotting with the indicated antibodies. (j) Relative mRNA levels of *NQO1* and *HMOX1* in Caki-2 sg*TSC1* cells treated with DMSO (−) or 25 nM rapamycin (+) overnight, measured by qRT-PCR. Data in (d), (f), (h), and (j) are presented as mean ± SD. \**P* < 0.05, \*\**P* < 0.01, \*\*\**P* < 0.001, and \*\*\*\**P* < 0.0001 (Student’s *t*-test).

Although the mechanisms by which increased ROS may enhance *NFE2L2* transcription remain incompletely understood, ROS and mTORC1 can activate NRF2 through well-characterized mechanisms involving protein stabilization. Central to this regulation is KEAP1, the substrate-recognition component of the CUL3–KEAP1 E3 ubiquitin ligase complex that targets NRF2 for proteasomal degradation^38^. ROS can oxidize cysteine residues in KEAP1, including C151, inducing conformational changes that impair NRF2 degradation. NRF2 can also be activated through SQSTM1/p62. mTORC1-mediated phosphorylation of human p62 at S349 (equivalent to mouse S351) enhances its binding to KEAP1, thereby competing with NRF2 for KEAP1 binding and promoting NRF2 stabilization^40,43^.

To test the hypothesis that increased mTORC1 signaling resulting from TSC1 loss can enhance NRF2 activity in human ccRCC, we used two *VHL^-/-^TSC1^+/+^*ccRCC cell lines, RCC4 and Caki-2. We generated CRISPR/Cas9-mediated *TSC1* knockout cells in both lines (RCC4 sg*TSC1* and Caki-2 sg*TSC1*). Loss of TSC1 increased NRF2 protein levels in both ccRCC cell lines and was accompanied by increased expression of the NRF2 target genes *NQO1* and *HMOX1* (Fig. 5c–f). As expected, TSC1 loss enhanced mTORC1 signaling, as indicated by increased phosphorylation of S6K at T389, 4EBP1 at T37/46, and p62 at S349 in both RCC4 sg*TSC1* and Caki-2 sg*TSC1* cells. Conversely, rapamycin treatment reduced NRF2 protein levels, suppressed *NQO1* and *HMOX1* expression, and decreased mTORC1-dependent phosphorylation in both TSC1-deficient cell lines (Fig. 5g–j).

### NRF2 is required for the growth of human *VHL^-/-^TSC1^-/-^* ccRCC cells

Human *VHL^-/-^TSC1^-/-^* tumors exhibit aggressive clinical features and can develop resistance to rapamycin analogs such as everolimus and temsirolimus, two FDA-approved therapies for ccRCC. As our studies thus far have implicated a critical role for NRF2 in the pathobiology of mouse *Vhl^-/-^Tsc1^-/-^* and human *VHL^-/-^TSC1^-/-^*ccRCC, we next explored the potential of targeting NRF2 signaling for the treatment of human kidney tumors lacking VHL and TSC1. To this end, we included HK-2, an immortalized human renal epithelial cell line, in our mechanistic studies. Combined CRISPR/Cas9-mediated knockout of *VHL* and *TSC1* in HK-2 cells resulted in increased NRF2 expression (Fig. 6a) and increased cellular proliferation in two independent *VHL/TSC1* double-knockout clones (Fig. 6b). Furthermore, siRNA-mediated knockdown of *NRF2* reduced the proliferation of HK-2 sg*VHL* sg*TSC1* cells to a greater extent than that of HK-2 sg*NT* cells, supporting a greater dependence of *VHL^-/-^TSC1^-/-^* cells on NRF2 for proliferation.

**Figure 6.**
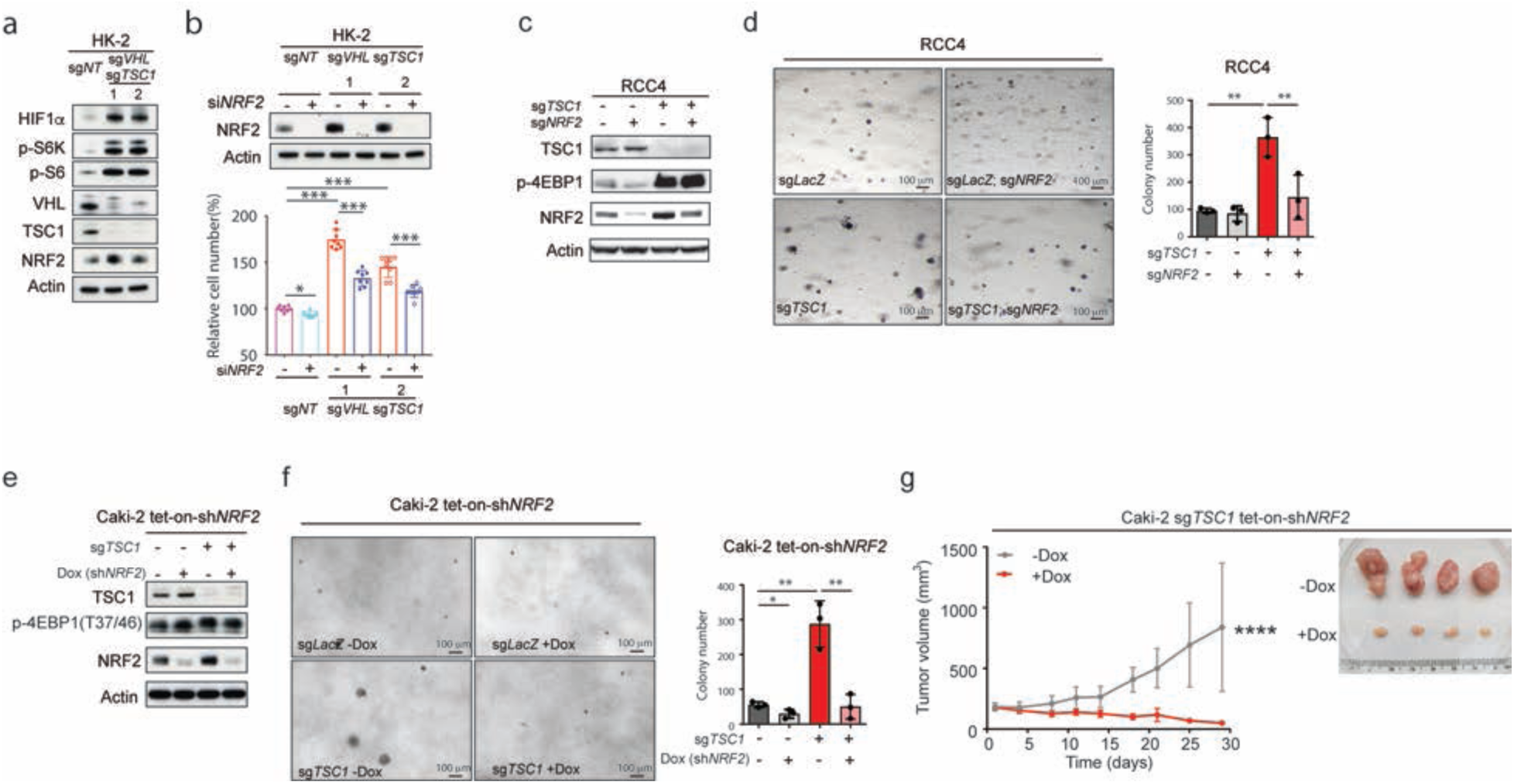
NRF2 is required for the growth of human *VHL^-/-^TSC1^-/-^* cells and tumors. (a) HK-2 cells expressing either sg*NT* or sg*VHL* sg*TSC1* were analyzed by immunoblotting with the indicated antibodies. (b) HK-2 cells expressing sg*NT* or sg*VHL* sg*TSC1* were transfected with si*NT* (−) or si*NRF2* (+) for 3 days. *NRF2* knockdown was assessed by immunoblotting, and relative cell proliferation was measured by the CellTiter-Glo assay. (c) RCC4 sg*LacZ* (−) and sg*TSC1* (+) cells expressing sg*LacZ* (−) or sg*NRF2* (+) were analyzed by immunoblotting with the indicated antibodies. (d) Representative images and quantification of soft-agar colony formation by RCC4 sg*LacZ* and sg*TSC1* cells expressing sg*LacZ* (−) or sg*NRF2* (+). (e) Caki-2 tet-on-sh*NRF2* cells expressing sg*LacZ* (−) or sg*TSC1* (+) were treated with doxycycline (Dox; 0 or 2 μg/ml) for 1 day and analyzed by immunoblotting with the indicated antibodies. (f) Representative images and quantification of soft-agar colony formation by Caki-2 tet-on-sh*NRF2* cells expressing sg*LacZ* or sg*TSC1* and treated with Dox (0 or 2 μg/ml). (g) Caki-2 sg*TSC1* tet-on-sh*NRF2* cells were implanted subcutaneously into the flanks of 8-week-old female NSG mice. When tumors reached approximately 100 mm³, mice were randomized to receive vehicle or 2 mg/ml doxycycline to induce sh*NRF2* (*n* = 8 per group). Tumor volumes were monitored for 28 days. Representative images of dissected tumors are shown. Data in (b), (d), (f), and (g) are presented as mean ± SD. Statistical significance in (b), (d), and (f) was determined by Student’s *t*-test and in (g) by two-way ANOVA. \**P* < 0.05, \*\**P* < 0.01, \*\*\**P* < 0.001, and \*\*\*\**P* < 0.0001.

We next turned to RCC4 and Caki-2 cells and compared their soft-agar colony-forming capacities according to their *TSC1* and *NRF2* status (Fig. 6c–f). Notably, knockout of *TSC1* in RCC4 cells resulted in an approximately threefold increase in soft-agar colony numbers, and this growth advantage was largely lost upon *NRF2* knockout (Fig. 6c,d). Similarly, knockout of *TSC1* in Caki-2 cells resulted in an approximately sixfold increase in soft-agar colony numbers, and this growth advantage was almost completely lost upon *NRF2* knockdown (Fig. 6e,f). Of note, NRF2 deficiency in parental RCC4 and Caki-2 cells also impaired colony-forming capacity, but to a lesser extent. We further performed tumor xenograft experiments. Remarkably, continuous regression of Caki-2 xenograft tumors was observed following induced *NRF2* knockdown (Fig. 6g). Altogether, these data identify NRF2 as a potential therapeutic target for human *VHL^-/-^TSC1^-/-^* ccRCC.

## Discussion

Our prior study of the mouse *Vhl^-/-^Pbrm1^-/-^* ccRCC model demonstrated that *Vhl^K-/-^Pbrm1^K-/-^* mice required ∼10 months to develop ccRCC and that mTORC1 activation played a critical role as a third oncogenic driver^9^. Here, prompted by genetic studies of human ccRCC, we investigated the *Vhl^K-/-^Tsc1^K-/-^* mouse model and unexpectedly found that these mice developed ccRCC as early as 7 weeks after birth, with 100% penetrance by ∼12 weeks, recapitulating the aggressive nature of its human counterpart. These findings indicate that two potent oncogenic signaling hubs, HIF and mTORC1, when simultaneously dysregulated, can cooperate to rapidly transform normal renal epithelium into aggressive ccRCC. Despite the potent oncogenic cooperation between VHL and TSC1 loss, biallelic *TSC1* loss is less frequent than inactivation of *VHL* (∼80%), *PBRM1* (∼40%), *BAP1* (∼10%), and *SETD2* (∼20%) in human ccRCC. The latter four genes are all located on chromosome 3p21–25, and loss of one copy of chromosome 3p, observed in >90% of ccRCC, constitutes the first oncogenic driver event in the pathogenesis of ccRCC^5^.

Our integrated mechanistic studies shed light on how these two oncogenic hubs reinforce each other to accelerate ccRCC development, likely through extensive transcriptional and metabolic crosstalk between HIF and mTORC1. mTORC1 is known to enhance HIF1α translation and thereby increase HIF signaling, whereas HIF1 activates REDD1, which in turn inhibits mTORC1 through the TSC1/2 complex, establishing a negative feedback loop. Thus, although loss of TSC1 can enhance HIF1α production through unrestrained mTORC1 signaling, HIF1α remains subject to VHL-mediated degradation. In the combined absence of VHL and TSC1, however, HIF activity is markedly enhanced, resulting in increased glycolytic metabolism at the expense of mitochondrial metabolism. At the same time, unrestrained mTORC1 activation appears to further reshape this metabolic and redox state by activating NRF2 signaling.

NRF2, the master regulator of the antioxidant response, also plays a broad role in intermediary metabolism by inhibiting lipogenesis, enhancing β-oxidation, facilitating flux through the pentose phosphate pathway, increasing NADPH regeneration, supporting purine biosynthesis, modulating mRNA translation, and activating macroautophagy^41,44,45^. NRF2 signaling was more strongly activated in *Vhl^K-/-^Tsc1^K-/-^* cortices than in WT, *Vhl^K-/-^*, or *Tsc1^K-/-^* cortices, likely reflecting the combined effects of increased ROS and unrestrained mTORC1 signaling. Although NRF2 has been implicated in the pathogenesis of papillary RCC, its involvement in *VHL^-/-^TSC1^-/-^*ccRCC was unexpected. Our integrated transcriptomic and metabolomic analyses identified prominent NRF2 pathway activation and redox and metabolic reprogramming in both mouse *Vhl^-/-^Tsc1^-/-^* and human *VHL^-/-^TSC1^-/-^* ccRCC models. Consistent with a functional requirement for this pathway, genetic silencing or deletion of NRF2 inhibited tumor growth in both *in vitro* cell-based and *in vivo* xenograft models, identifying NRF2 as a critical dependency and potential therapeutic vulnerability in *VHL^-/-^TSC1^-/-^* ccRCC.

Beyond NRF2 dependency, our metabolomic studies also implicated heightened glutaminolysis (Fig. 3d) as an additional metabolic vulnerability. Glutaminolysis generates α-ketoglutarate (αKG) to support TCA-cycle anaplerosis^46^ and glutamate for glutathione synthesis, thereby linking glutamine metabolism to both bioenergetic and redox demands. Furthermore, elevated glutamine levels in *Vhl^K-/-^Tsc1^K-/-^* renal epithelium may contribute to the further increase in mTORC1 signaling observed in *Vhl^K-/-^Tsc1^K-/-^* compared with *Tsc1^K-/-^* cortices, as glutamine can promote mTORC1 activation by facilitating its recruitment to the lysosomal surface^47^. Thus, glutaminase inhibition may provide an additional therapeutic strategy for ccRCC. Indeed, our clinical trial (NCT03163667) combining the mTORC1 inhibitor everolimus with the glutaminase inhibitor CB-839 demonstrated greater clinical activity than everolimus alone in ccRCC^48^. Together, these findings support further exploration of NRF2-dependent antioxidant pathways and glutamine metabolism as therapeutic targets in molecularly defined subsets of ccRCC^49^.

## Methods

### Mice

*Ksp-Cre* transgenic mice^31^ (B6.Cg-Tg(Cdh16-cre)91Igr/J; RRID:IMSR_JAX:012237), *Vhl^F/F^*mice^50^ (B6.129S4(C)-Vhltm1Jae/J; RRID:IMSR_JAX:012933), and *Tsc1^F/F^* mice^51^ (STOCK Tsc1tm1Djk/J; RRID:IMSR_JAX:005680) were obtained from The Jackson Laboratory. These mouse strains were bred to generate *Vhl^F/F^Ksp-Cre*^+^ (*Vhl^K-/-^*)*, Tsc1^F/F^Ksp-Cre^+^* (*Tsc1^K-/-^*), and *Vhl^F/F^Tsc1^F/F^Ksp-Cre^+^* (*Vhl^K-/-^Tsc1^K-/-^*) mice. All mice were maintained on a mixed B6/129 genetic background. NSG mice (NOD.Cg-Prkdc^scid^ Il2rg^tm1Wjl^/SzJ) used for xenograft studies were also obtained from The Jackson Laboratory. All mouse experiments were performed under a protocol approved by the Institutional Animal Care and Use Committee (IACUC) at Memorial Sloan Kettering Cancer Center (MSKCC). All animal work was performed in accordance with NIH guidelines (Guide for the Care and Use of Laboratory Animals, National Academies Press, 2011) and in compliance with MSKCC institutional requirements.

### Human tumor samples

FFPE tissue samples were obtained from primary nephrectomy specimens collected at MSKCC under institutional review board–approved tissue collection protocols, with informed consent from all patients. All cases were reviewed by genitourinary pathologists and confirmed to meet the diagnostic criteria for renal cell carcinoma based on the current World Health Organization classification and consensus criteria of the International Society of Urological Pathology.

### Cell lines and treatments

Human proximal tubular epithelial HK-2 cells (CRL-2190) and human clear cell renal cell carcinoma Caki-2 cells (HTB-47) were obtained from ATCC and cultured according to the manufacturer’s recommendations. Human clear cell renal cell carcinoma RCC4 cells (CVCL_0498) were obtained from the Cancer Cell Line Encyclopedia (CCLE). For rapamycin treatment experiments, RCC4 sg*TSC1* and Caki-2 sg*TSC1* cells were treated with 25 nM rapamycin (S1039, Selleckchem) or DMSO overnight before analysis by immunoblotting and qRT-PCR.

### Mouse MRI

Mouse MRI scans were performed using either a 4.7-T or 7-T Bruker BioSpec scanner (200 or 300 MHz; Bruker BioSpin MRI GmbH, Ettlingen, Germany) equipped with 640 mT/m, 115-mm-ID or 300 mT/m, 200-mm-ID gradients, respectively (Resonance Research, Inc., Billerica, MA). RF excitation and signal acquisition were performed using a custom-built quadrature birdcage resonator with a 32-mm inner diameter (Stark Contrast MRI Coils Research Inc., Erlangen, Germany). Mice were anesthetized with 1% isoflurane (Baxter Healthcare Corp., Deerfield, IL) in oxygen, and respiration was monitored using a small-animal physiological monitoring system (SA Instruments, Inc., Stony Brook, NY). Scout images in three orthogonal orientations were first acquired for animal positioning. For kidney imaging, coronal T2-weighted images were acquired using a fast spin-echo RARE (Rapid Acquisition with Relaxation Enhancement) sequence with TR = 1.5 s, TE = 48 ms, RARE factor = 8, slice thickness = 0.7 mm, field of view = 30 mm, in-plane resolution = 117 × 156 μm, and 18 averages.

### Serum creatinine measurement

Blood was collected from mice by retro-orbital sampling. Serum creatinine levels were measured by the Laboratory of Comparative Pathology (LCP) at Memorial Sloan Kettering Cancer Center using a Beckman Coulter AU680 analyzer.

### Histopathology

Mouse kidneys were harvested and fixed overnight in 10% (vol/vol) neutral-buffered formalin. Fixed kidneys were then processed through ethanol and xylene, embedded in paraffin, sectioned at 5 µm, and stained with hematoxylin and eosin (H&E). Selected slides were reviewed by board-certified pathologists.

### Immunohistochemistry

Paraffin-embedded tissues were sectioned at 5 µm and stained by the Pathology Core at MSKCC. Immunohistochemistry (IHC) was performed using the Ventana Discovery XT platform (Ventana Medical Systems). Briefly, formalin-fixed, paraffin-embedded tissues were sectioned at 5 µm and air-dried overnight. Sections were deparaffinized, rehydrated, and subjected to heat-induced epitope retrieval (HIER), followed by incubation with the indicated primary antibodies. Slides were subjected to polymer treatment to reduce background staining in kidney tissue. For signal detection, OmniMap anti-Rb DAB HRP was used according to the manufacturer’s instructions. Slides were counterstained with hematoxylin. IHC for 2SC was performed as described previously^52^. Appropriate positive and negative controls were included in each IHC run. IHC for Ki-67 (HIER at pH 9.0) and cleaved caspase-3 (HIER at pH 6.0) was performed by the Laboratory of Comparative Pathology (LCP) at MSKCC using a Leica Bond RX automated staining platform with a polymer detection system (Novocastra Bond Polymer Refine Detection, Leica Biosystems) and hematoxylin counterstaining. TUNEL staining was performed by LCP at MSKCC as described previously^53^. Fluorescein-conjugated lotus tetragonolobus lectin (LTL) staining (FL-1321, Vector Laboratories) was performed on rehydrated slides after antigen retrieval for 4 h at room temperature, followed by DAPI counterstaining and Sudan Black treatment. PAS/PAS-diastase (PASD) and Oil Red O staining were performed using standard methods^54,55^. Frozen tissue sections were used for Oil Red O staining, whereas fixed paraffin-embedded tissue sections were used for PAS and PASD staining. Antibodies used for immunohistochemistry included anti-carbonic anhydrase IX (NB100-417, Novus), anti-phospho-4E-BP1 (Thr37/Thr46) (2855, Cell Signaling Technology), anti-phospho-S6 ribosomal protein (Ser240/244) (5364, Cell Signaling Technology), anti-Tamm-Horsfall glycoprotein (THP) (J65429, Alfa Aesar/Thermo Fisher Scientific), anti-phospho-AKT (Ser473) (4060, Cell Signaling Technology), anti-CD31 (M0823, Agilent), anti-phospho-p44/42 MAPK (ERK1/2) (Thr202/Tyr204) (4370, Cell Signaling Technology), anti-NFE2L2 (ab62352, Abcam), anti-HIF-1α (10006421, Cayman Chemical), anti-8-hydroxy-2′-deoxyguanosine (ab48508, Abcam), anti-Ki-67 (ab16667, Abcam), and anti-cleaved caspase-3 (Asp175) (9661, Cell Signaling Technology).

### RNA isolation, RNA-seq, and analysis

RNA was isolated using TRIzol (15596018, Thermo Fisher Scientific) and purified using the RNeasy Mini Kit (74104, Qiagen). RNA samples were prepared from 4-week-old *Ksp-Cre* (WT), *Vhl^K-/-^*, *Tsc1^K-/-^*, and *Vhl^K-/-^Tsc1^K-/-^* mice. Total RNA was processed by the Integrated Genomics Operation (IGO) at MSKCC using the TruSeq RNA Sample Prep Kit according to the manufacturer’s recommendations. Briefly, cDNA libraries were prepared from poly(A)-selected RNA by end repair, A-tailing, adaptor ligation, and PCR enrichment. Libraries were sequenced on an Illumina HiSeq 2500 platform using 50-bp paired-end reads, with approximately 40 million reads per sample. Sequencing reads were mapped to the reference mouse genome, quantified, and analyzed for differential expression using Partek Flow. Genes with an FDR < 0.05 were used for clustering and pathway analyses. Hierarchical agglomerative clustering was performed and visualized using the pheatmap package in R. Gene Ontology (GO) analysis of the mouse RNA-seq data was performed using ClueGO as previously described^9^. Gene set enrichment analysis (GSEA) was performed using default settings with the MSigDB H collection as the gene set database. Each gene set was permuted 1,000 times to calculate *P* and FDR values. Correlation analyses of gene signatures in the TCGA and CCLE cohorts shown in Fig. 5 were performed using the R2 Genomics Analysis and Visualization Platform (https://hgserver1.amc.nl/cgi-bin/r2/main.cgi).

### Reverse transcription and quantitative PCR

Total RNA was isolated from 4-week-old *Ksp-Cre* (WT), *Vhl^K-/-^, Tsc1^K-/-^,* and *Vhl^K-/-^Tsc1^K-/-^* mice or from human cell lines as described above. Reverse transcription was performed using oligo(dT) primers (AM5730G, Thermo Fisher Scientific) and random decamer primers (AM5722G, Thermo Fisher Scientific) with SuperScript II (18064014, Thermo Fisher Scientific). Quantitative PCR was performed in duplicate using SYBR Green Master Mix (4309155, Thermo Fisher Scientific) and the indicated gene-specific primers on a ViiA™ 7 Real-Time PCR System (Applied Biosystems). Gene-specific primers are listed in Supplementary Table 7. Data were analyzed as described previously^9^ by normalization to *Actb*.

### Metabolomics analysis

For metabolomics analysis, renal cortices from 4-week-old *Ksp-Cre* (WT), *Vhl^K-/-^, Tsc1^K-/-^,* and *Vhl^K-/-^Tsc1^K-/-^*mice were dissected, immediately snap-frozen, and stored at −80°C until analysis. Samples and standards were analyzed as described previously^36^. Briefly, dried extracts were dissolved in 10 µl of 40 mg/ml O-methoxyamine hydrochloride in pyridine and shaken at 30°C for 1.5 h. Ninety microliters of N-methyl-N-(trimethylsilyl)-trifluoroacetamide containing fatty acid methyl ester retention-index markers was then added to each tube, followed by shaking at 37°C for 30 min. Within 48 h of derivatization, all samples were injected onto an Agilent 6890 gas chromatograph coupled to a LECO Pegasus III time-of-flight mass spectrometer. A Restek RTX-5Sil MS column (95% dimethyl/5% diphenyl polysiloxane; 30 m length, 0.25 mm inner diameter, 0.25 µm film thickness) with a 10 m guard column was used. Data were acquired over a mass range of *m/z* 85–500 at 17 spectra/s with a detector voltage of 1,850 V. Peaks were deconvoluted and detected using LECO ChromaTOF software and matched to the FiehnLib mass spectral and retention-time library. BinBase software was used for post-curation and peak replacement. Data were normalized using the summed intensities of all known compounds. Principal component analysis, hierarchical clustering, and quantitative enrichment analysis of metabolites across samples were performed using MetaboAnalyst 3.0 (https://www.metaboanalyst.ca/).

### GSH and GSSG quantitation

GSH and GSSG levels were measured in mouse kidney tissue homogenates using the GSH/GSSG-Glo Assay Kit (Promega) as described previously^56^. Briefly, frozen tissues were homogenized in 5% sulfosalicylic acid, and proteins were precipitated by centrifugation. Acid extracts were neutralized with 0.5 mol/L HEPES (pH 8.0) and, when necessary, diluted with 0.25 mol/L HEPES (pH 7.5). GSH and GSSG standard curves were prepared using the same concentrations of sulfosalicylic acid and HEPES buffers used for tissue extract preparation and neutralization.

### CRISPR knockout

sgRNA and Cas9 were encoded in the lentiviral vector lentiCRISPRv2 (Addgene). CRISPR sgRNA sequences were as follows: sg*VHL*, GAGTCCGGCCCGGAAGAGTC; sg*TSC1*, TTTATCCATCCTCTCGTTAC; sg*NT*, TGCGAATACGCCCACGCGAT; and sg*NRF2*, CATACCGTCTAAATCAACAG. Lentiviruses were produced in 293T cells by co-transfection with the packaging plasmids psPAX2 and VSV-G. Cells were infected with lentiviruses for 12 h and selected with 2 µg/ml puromycin for 1 week before further analysis.

### Immunoblot analysis

Immunoblot analysis was done as described previously^10^. Briefly, membranes were blocked in 5% nonfat dry milk for 1 h at room temperature and incubated with the indicated primary antibodies. Membranes were then incubated with horseradish peroxidase–conjugated anti-mouse or anti-rabbit IgG secondary antibodies, and signals were detected using enhanced chemiluminescence. Antibodies used for immunoblotting included anti-NFE2L2 (ab62352, Abcam), anti-phospho-p70 S6 kinase (Thr389) (9205, Cell Signaling Technology), anti-p70 S6 kinase (9202, Cell Signaling Technology), anti-phospho-4E-BP1 (Thr37/Thr46) (2855, Cell Signaling Technology), anti-4E-BP1 (9452, Cell Signaling Technology), anti-phospho-SQSTM1/p62 (Ser349) (95697, Cell Signaling Technology), anti-SQSTM1/p62 (ab56416, Abcam), anti-hamartin/TSC1 (6935, Cell Signaling Technology), anti-β-actin (4970, Cell Signaling Technology), anti-vinculin (4650, Cell Signaling Technology), anti-HIF-1α (10006421, Cayman Chemical), anti-VHL (2738, Cell Signaling Technology), and anti-phospho-S6 ribosomal protein (Ser240/244) (5364, Cell Signaling Technology).

### siRNA, shRNA, and cell proliferation assays

HK-2 sg*NT* or sg*VHL* sg*TSC1* cells were transfected with control or *NRF2* (*NFE2L2*) siRNA (Dharmacon; si*Control*: ON-TARGETplus Non-targeting Pool siRNA, D-001810-10-05; si*NFE2L2*: ON-TARGETplus NFE2L2 Pool siRNA, L-003755-00-0005) using Lipofectamine RNAiMAX (Thermo Fisher Scientific) according to the manufacturer’s instructions. For cell proliferation assays, siRNA-transfected cells were seeded into 96-well plates at 1,000 cells per well. Cell proliferation was quantified using the CellTiter-Glo assay (Promega). Tet-on shRNA lentiviruses were produced in 293T cells by co-transfection with the packaging plasmids pCMVΔR8.2 and VSV-G. Cells were infected with lentiviruses for 12 h and selected with 2 µg/ml puromycin for 1 week before further analysis.

### Soft agar assay and quantification

Soft agar was prepared using Noble agar and the corresponding cell culture medium. Bottom layers containing 0.6–1% agar in culture medium were allowed to solidify, after which cells suspended in 0.3–0.4% agar in culture medium were overlaid as the top layer. Once the top layer had solidified, complete culture medium was added and replenished every 3 days. To induce Tet-on construct expression, doxycycline was added to the medium 12 h after the soft agar had solidified. When colonies in control cultures reached a diameter greater than 100 μm, colonies were counted using a GelCount colony counter (Oxford Optronix), and microscopic images were acquired at low magnification using a Nikon Eclipse Ti microscope. The assay was performed in triplicate.

### Xenograft studies

For subcutaneous xenograft studies, 5 × 10^6^ Caki-2 sg*TSC1* cells expressing the Tet-on sh*NRF2* construct were suspended in Matrigel (356234, BD Biosciences) and injected subcutaneously into both flanks of 8-week-old female NSG mice. Mice received doxycycline (2 mg/ml; Sigma-Aldrich) and sucrose (50 mg/ml) in the drinking water to induce sh*NRF2* expression. Tumor dimensions were measured with calipers, and tumor volume was calculated as ½ × width² × length.

### Statistical analysis

Statistical analyses were performed using GraphPad Prism (GraphPad Software) unless otherwise indicated. Survival differences were evaluated using the log-rank (Mantel–Cox) test. Kidney weights and serum creatinine levels were compared using the Mann–Whitney test. Comparisons of qRT-PCR measurements, GSH/GSSG measurements, cell proliferation, and soft-agar colony formation were performed using Student’s *t*-test, and data are presented as mean ± SD. Xenograft tumor growth curves were analyzed by two-way ANOVA. For RNA-seq analyses, differentially expressed genes were identified using Benjamini–Hochberg-adjusted *P* values, with an FDR cutoff of 0.05. Gene set enrichment analysis (GSEA) was performed using 1,000 permutations to calculate *P* and FDR values. For global metabolomic analyses, differences among the four genotypes were assessed by one-way ANOVA with FDR correction, whereas pairwise comparisons of individual metabolite levels shown in the figures were performed using Student’s *t*-test. *P* < 0.05 was considered statistically significant unless otherwise indicated. Statistical significance is denoted as \**P* < 0.05, \*\**P* < 0.01, \*\*\**P* < 0.001, and \*\*\*\**P* < 0.0001.

## Data availability

Raw RNA-seq data have been deposited in the Gene Expression Omnibus (GEO) under accession number GSE131735. Normalized RNA-seq gene counts are available in Supplementary Table 5. Metabolomics data are available in Supplementary Table 6.

## Acknowledgements

This work was supported by the NIH grant (R01 CA125562) to E. Cheng. This work was also supported by the NCI Cancer Center Support Grant (P30CA008748).

## Author contributions

J.X. and X.C. designed and conducted experiments, and analyzed data. J.J.H. and E.H.C. designed research and supervised the project. O.A.A., Y.B.C., A.B., T.O., and S.T. conducted some experiments. E.R. analyzed some data. Y.B.C. and S.K.T. performed histological assessment. R.H.W. supervised some experiments.

---

**Supplementary Fig. 1.**
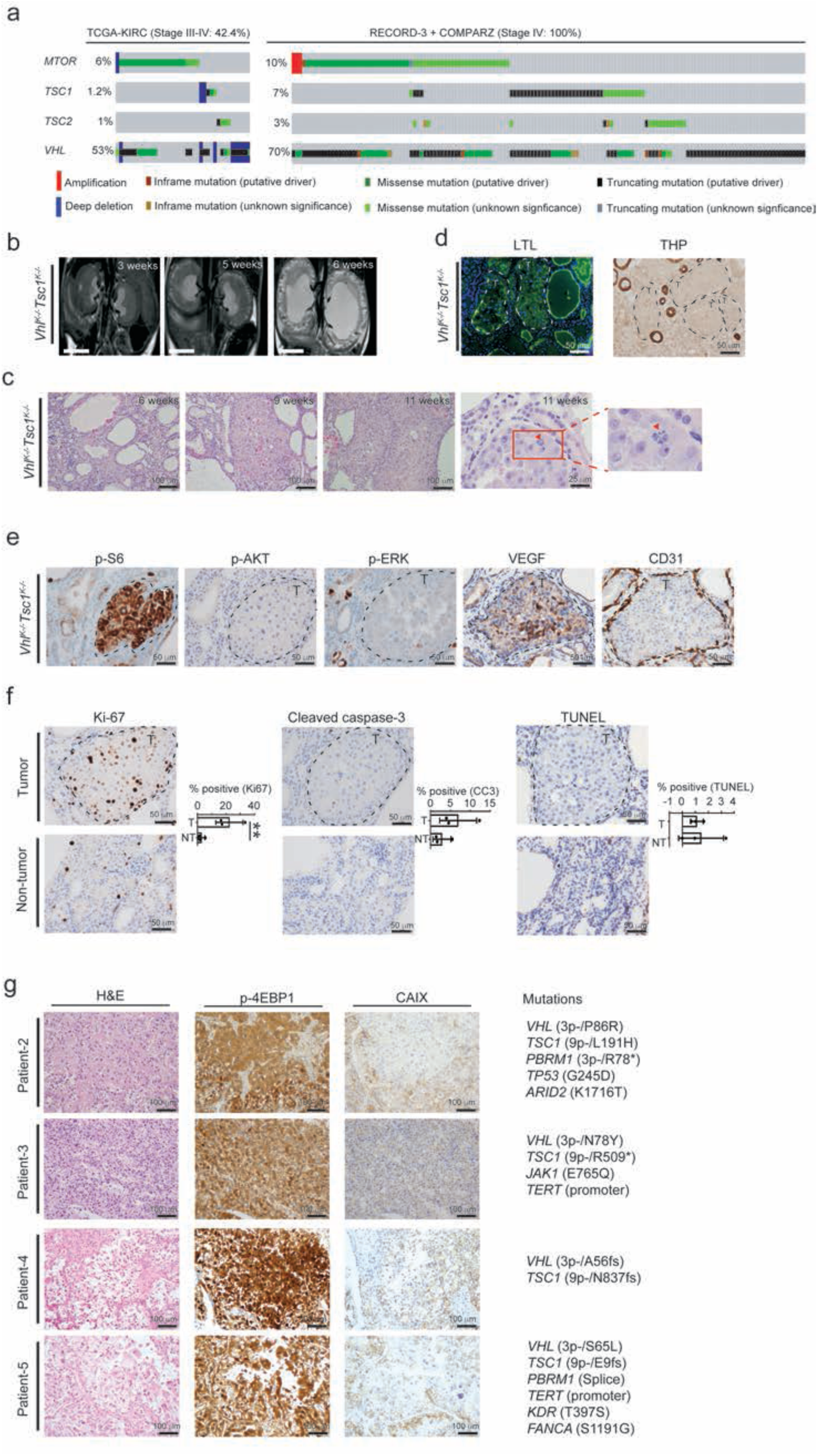
Oncoprint of human ccRCC and morphological and immunohistochemical analyses of *Vhl^K-/-^Tsc1^K-/-^*mouse kidneys and human ccRCC. (a) Oncoprint of mutation profiles of selected genes from the TCGA-KIRC, RECORD-3, and COMPARZ studies. Only a subset of *VHL* mutations is shown. (b) MRI images of kidneys from *Vhl^K-/-^Tsc1^K-/-^* mice at the indicated ages. (c) Representative H&E-stained kidney sections from *Vhl^K-/-^Tsc1^K-/-^*mice at the indicated ages. The red arrow indicates an example of a mitotic figure in a *Vhl^-/-^Tsc1^-/-^* kidney tumor from an 11-week-old mouse. (d) Representative lotus tetragonolobus lectin (LTL; a proximal convoluted tubule marker) and Tamm-Horsfall protein (THP; a distal tubule marker) staining of tumor regions (denoted as T) in kidneys from 11-week-old *Vhl^K-/-^Tsc1^K-/-^*mice. (e) Representative immunohistochemical staining for p-S6, p-AKT, p-ERK, VEGF, and CD31 in tumor regions (denoted as T) of kidneys from 11-week-old *Vhl^K-/-^Tsc1^K-/-^* mice. (f) Representative immunohistochemical staining for Ki-67 and cleaved caspase-3 and TUNEL staining in tumor (T) and non-tumor (NT) regions of kidneys from 11-week-old *Vhl^K-/-^Tsc1^K-/-^* mice. Quantification of Ki-67, cleaved caspase-3, and TUNEL staining is shown. Three microscopic fields were randomly selected, and the percentage of positively stained cells relative to the total number of cells was calculated. Data are shown as the mean from kidney sections of three different mice. Data are presented as mean ± SD. \*\**P* < 0.01 (Student’s *t*-test). (g) Representative H&E-stained sections and immunohistochemical staining for p-4EBP1 and CAIX in human renal tumors with *VHL* and *TSC1* loss. Mutation profiles determined by MSK-IMPACT analysis are indicated.

**Supplementary Fig. 2.**
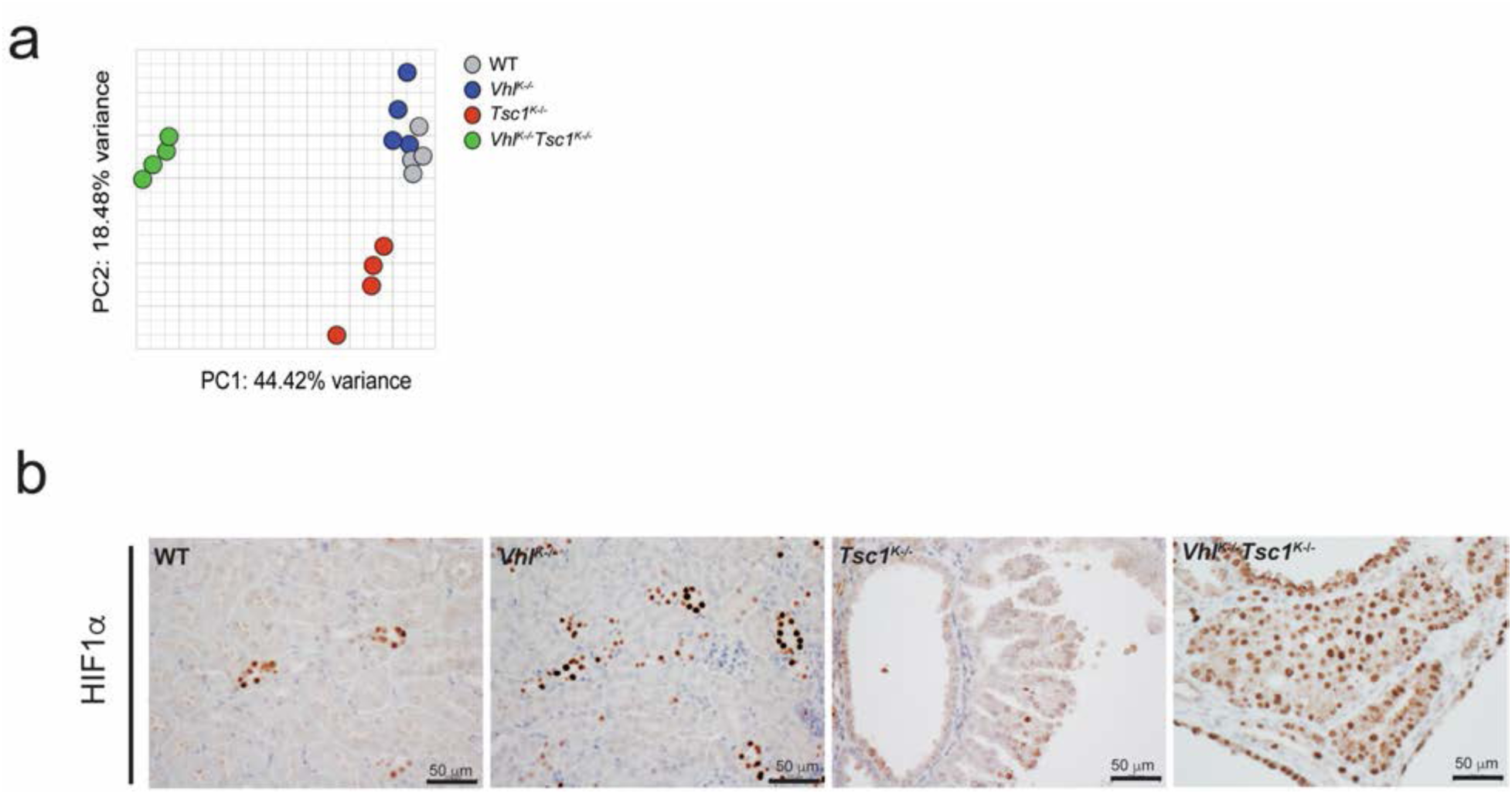
Additional transcriptomic and immunohistochemical analyses. (a) Principal component analysis of RNA-sequencing data from renal cortices of 4-week-old WT, *Vhl^K-/-^, Tsc1^K-/-^,* and *Vhl^K-/-^Tsc1^K-/-^* mice. (b) Representative immunohistochemical staining for HIF1α in renal cortices from 11-week-old WT, *Vhl^K-/-^, Tsc1^K-/-^,* and *Vhl^K-/-^Tsc1^K-/-^*mice.

**Supplementary Fig. 3.**
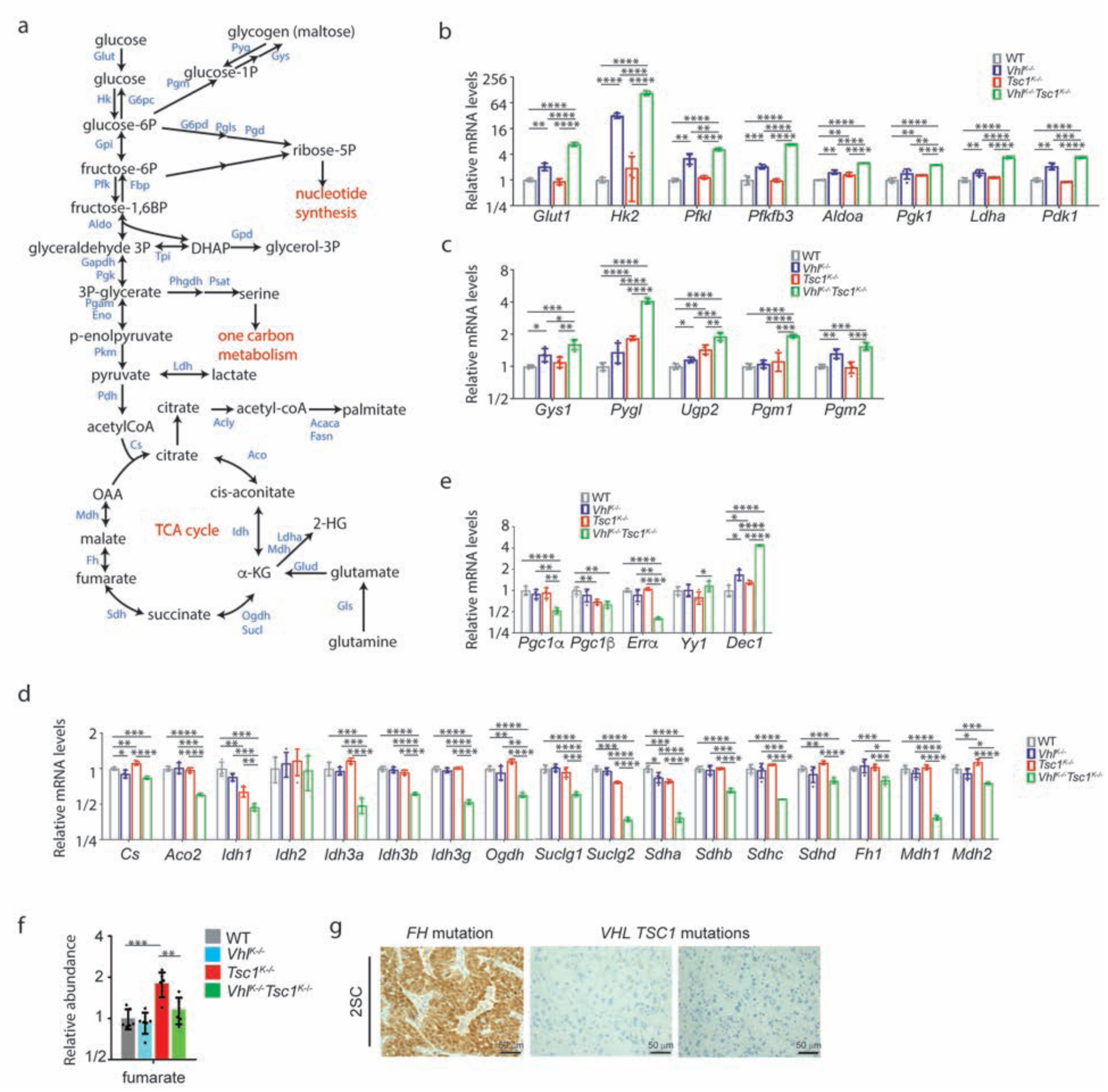
Additional transcriptomic, metabolomic, and immunohistochemical analyses of metabolic pathways. (a) Schematic of glucose and glutamine metabolism. (b) Relative mRNA levels of genes involved in glycolysis in renal cortices from 4-week-old mice of the indicated genotypes. (c) Relative mRNA levels of genes involved in glycogen metabolism in renal cortices from 4-week-old mice of the indicated genotypes. (d) Relative mRNA levels of genes involved in the TCA cycle in renal cortices from 4-week-old mice of the indicated genotypes. (e) Relative mRNA levels of genes regulating mitochondrial biogenesis in renal cortices from 4-week-old mice of the indicated genotypes. (f) Relative abundance of fumarate in renal cortices from 4-week-old mice of the indicated genotypes. (g) Representative immunohistochemical staining for S-(2-succinyl)cysteine (2SC) in human renal tumors from patients 2 and 3 in Supplementary Fig. 1g and in an FH-mutant renal tumor used as a positive control. Data in (b–f) are presented as mean ± SD. \**P* < 0.05, \*\**P* < 0.01, \*\*\**P* < 0.001, and \*\*\*\**P* < 0.0001 (Student’s *t*-test).

## Supplementary Table Legends

**Supplementary Table 1.** Mouse information and renal histopathologic findings.

**Supplementary Table 2.** Clinicopathologic and genomic features of human renal tumors with *VHL* and *TSC1* mutations.

**Supplementary Table 3.** Gene Ontology and KEGG pathways enriched in Clusters I–III identified by ClueGO analysis.

**Supplementary Table 4.** GSEA of the “Signal transduction” and “Regulation of gene expression” groups within Cluster II identified in Fig. 2a,b.

**Supplementary Table 5.** Normalized RNA-seq gene counts in renal cortices from 4-week-old WT, *Vhl^K-/-^*, *Tsc1^K-/-^*, and *Vhl^K-/-^Tsc1^K-/-^* mice.

**Supplementary Table 6.** mTIC-normalized metabolite abundances in renal cortices from 4-week-old WT, *Vhl^K-/-^*, *Tsc1^K-/-^*, and *Vhl^K-/-^Tsc1^K-/-^* mice.

**Supplementary Table 7.** Primer sequences used for qRT-PCR.

